# Noise-robust, physical microscopic deconvolution algorithm enabled by multi-resolution analysis regularization

**DOI:** 10.1101/2023.03.02.530744

**Authors:** Yiwei Hou, Wenyi Wang, Yunzhe Fu, Xichuan Ge, Meiqi Li, Peng Xi

## Abstract

Despite the grand advances in fluorescence microscopy, the photon budget of fluorescent molecules remains the fundamental limiting factor for major imaging parameters, such as temporal resolution, duration, contrast, and even spatial resolution. Computational methods can strategically utilize the fluorescence photons against the imaging noise, to break the abovementioned limits. Here, we propose a multi-resolution analysis (MRA) approach to recharacterize and extract the two main characteristics of fluorescence images: (1) high contrast across the edge, and (2) high continuity along the edge. By regularizing the solution using framelet and curvelet domain sparsity, we develop MRA deconvolution algorithm for fluorescence image, which allows fine detail recovery even with negative signal-to-noise-ratio (SNR), and can provide more than 2-fold physical resolution enhancement with conspicuously fewer artifacts than maximum likelihood estimation (MLE) methods. Furthermore, we develop DeepMRA deconvolution algorithm that can provide computational background inhibition through a bias thresholding mechanism while deconvolving a fluorescence image. Compared with conventional background mitigation schemes, this novel deconvolution canonical form can deal with severer background and better preserve the high-frequency and low-intensity details, which are commonly disrupted by other algorithms. We demonstrate that the MRA and DeepMRA deconvolution algorithms can improve the SNR and resolution of biological images in various microscopies, such as wide-field, confocal, spinning-disk confocal (SD-confocal), light-sheet, structured illumination microscopy (SIM), and stimulated excitation depletion (STED) microscopy.

## Introduction

Fluorescence microscopy lays the foundation of modern optical microscopy in the application of live-cell imaging^1^. The birth of fluorescence super-resolution (SR) microscopy^2–5^ even breaks the diffraction limit of conventional optical microscopy, which arises from the wave nature of the illumination and detection signals. Although in principle the spatial resolution of SR can reach single-molecule resolution, in practice it is limited by the detectable fluorescent photon flux: higher spatial resolution implies more pixels and fewer photons per pixel, and it holds true also for temporal resolution. On the one hand, the development of brighter and higher photostable dyes, with more sensitive and faster detection techniques are preferred^6, 7^. On the other hand, computational image processing techniques^8–17^ that can recover the fluorescence signal from the abundant noise are highly welcome as it paves a new avenue for further improving the spatiotemporal resolution and signal-to-noise-ratio (SNR) of fluorescence microscopy.

Deconvolution algorithms based on the image degeneration model are one of the most effective schemes to recover fluorescence images from optical blurring and noise. Classical inverse problem solvers such as Wiener filtering and Richardson-Lucy (RL) iteration work at moderate noise levels but are feeble for low-SNR images. Since the SNR of fluorescence images is generally poor, especially in ultrafast or long-term imaging, the noise-robustness feature is highly desired, which is achieved by the regularization-based deconvolution method. For example, using the continuity prior knowledge of fluorescence image, total-variation (TV)^8^ and Hessian^9^ regularization-based deconvolution algorithms were developed for low-SNR SIM image compensation. Improving the resolution of fluorescence images under noise conditions is even more challenging since high fidelity is required for life science research. Sparse deconvolution^10^ based on the numerical sparsity and sample continuity was proposed to enhance the conventional RL iteration, which can improve the spatial resolution up to 60 nm under SIM imaging mode. However, limited by their naïve assumptions about the fluorescence image, losses in spatial and temporal details are usually inevitable. Moreover, due to the artifact-proneness feature^18^, the fidelity of the statistical maximum likelihood estimation (MLE) method remains unclear. Another method of resolution improvement directly minimizes the fidelity penalty such as the fast iteration soft-thresholding algorithm (FISTA)^19^, which is a physical deconvolution approach since it mimics the process of diffraction elimination. However, it is easily affected by noise and is discordant with variance-based denoising regularization, thus not showing good performance in the real microscopic image. Besides the deconvolution method, recently some deep learning approaches^11–17^ have also been proposed to enhance fluorescence image in various microscopies. Yet, the caveat of the deep learning approaches remains: without proper training of the model, it is impossible to reach a trustworthy result, despite that the images are very appealing to human eye. Moreover, proper training often implies that at least similar phenomena were imaged previously, which is impossible for new discoveries.

As an improvement of the conventional data processing method, the wavelet transform theory was proposed by mathematicians in the early 20th century^20^, and was first applied in engineering for seismic wave data analysis until 1984^21^. Later, the theory of multi-resolution analysis (MRA)^22–24^ that decomposes the signal into a series of resolution levels was proposed, which provides a novel idea for signal space segmentation. The traditional orthogonal wavelet^25,26^ is the first developed MRA tool that achieves optimal results in a variety of image processing tasks^27–32^. As the importance of sparsity in signal processing was clarified^33^, some redundant multiscale bases^34–38^ that can provide a sparser representation of signals were further developed, which show superior performance^39–42^. However, possibly due to the distinct difference between macro-scale and fluorescence images, the MRA approach for fluorescence and SR image enhancement has been an underexplored domain so far.

In this work, we introduce the MRA approach to characterize the two major characteristics of fluorescence microscopic image: high contrast across the edge, and high continuity along the edge. This new regularization standard outperforms the most advanced variance-based regularization with conspicuously improved signal-noise discrimination ability, and even supports significant physical resolution improvement in the practical microscopic image. With MRA deconvolution, we have successfully reconstructed a series of cellular organelles and cytoskeletons including the actin filaments, microtubules, mitochondria, ER network, etc., with high fidelity at low SNR level. A 30-nm spatial resolution is attained with SIM imaging to the dual-line resolution target. To resolve the sample feature under strong background, we further present DeepMRA, which can mitigate background and provide computational sectioning in each deconvolution iteration, just like confocal microscopy. DeepMRA can better preserve high-frequency and low-intensity details at high surrounding local noise and background levels. The wide feasibility of MRA and DeepMRA is verified in main-stream fluorescence microscopies such as wide-field, confocal, spinning-disk confocal (SD-confocal) light-sheet, SIM, and stimulated excitation depletion (STED) microscopy.

## Results

### MRA deconvolution algorithm

Traditional deconvolution algorithm for fluorescence images is developed based on the spatial or Fourier domain understanding of the degeneration process (**Supplementary Note 1**). One of the most important backbones for these analytical models is the continuity prior knowledge of fluorescence image, which is characterized by the spatial derivative. However, this method tangles the continuity of the cell structure itself and its border with the background (**Supplementary Fig. 1**), and considering only the continuity feature is incomplete for describing a fluorescence image. These drawbacks fundamentally limit the performance of these conventional deconvolution algorithms. Here we sake to solve this dilemma by setting a new prior knowledge standard for fluorescence images in the multiscale base transform domain. We recharacterize the features of fluorescence images as: (1) high contrast across the edge, and (2) high continuity along the edge. High contrast sharp edge is an important characteristic of fluorescence images because the fluorophore specifically attaches a biological structure other than its surrounding background due to the superior specificity, which can be effectively detected by framelet^36^. Continuity is also an important prior knowledge of fluorescence images because of the connectivity of the biological specimen, and sufficient spatial sampling nature of fluorescence microscopes. Instead of the spatial derivative-based method, we employ curvelet^34,35^ to represent this feature because it specifically characterizes the along-edge continuity, which distinguishes the mix of biological structure and the background.

To verify the above analysis, we added Gaussian noise to the fluorescence image of various organelles and calculated the sparsity in the framelet and curvelet domain. The results show that both the framelet and curvelet domain sparsity decreases with the increase of noise level (**Supplementary Fig. 2**). Increasing the sparsity of the framelet and curvelet coefficients through hard thresholding, the sharp edge and along-edge continuity feature of the fluorescence image can be extracted from the noise contamination (**Supplementary Note 2** and **Supplementary Fig. 3**). Based on above assumptions and verifications, we propose the co-sparsity of the framelet and curvelet coefficients as the regularization for the fluorescence image deconvolution model, termed MRA deconvolution:

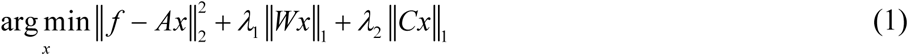

where *f* is the input degenerated image, *A* denotes the blur kernel in the matrix form, *x* is the recovered image, *W* denotes the linearized piece-wise framelet transform, *C* denotes the curvelet transform, *λ*1 and *λ*2 are two regularization parameters, ||1 and ||2 notions denote the *l*1 and *l*2 norm, respectively. The first term ensures the physical nature of the deconvolution process, which is also called the fidelity term. The second and third terms are the sparsity of the recovered image in the framelet and curvelet domain, respectively. FISTA is employed to minimize this optimization problem (**Supplementary Note 3.1**). A small number of RL iteration can be executed after the iteration to enhance the image contrast.

For live-cell imaging, the image fidelity would be inevitably damaged if a short exposure or low illumination intensity is used, which are usually essential for capturing the dynamic long-term cell movement. We expect the MRA deconvolution to recover information from ultralow-SNR images based on the strong signal-extracting capability of the framelet and curvelet transforms. Besides the spatial characteristics of fluorescence images, the continuity in an additional spatial direction and temporal domain can also be utilized to recover a low-SNR image stack. To enhance the ability of MRA for deconvolving time-lapse or 3D images with good continuity, we develop a spatiotemporal continuity denoising scheme based on the soft-thresholding of 3D framelet and 3D dual-tree complex wavelet (DTCW)^43,44^ coefficients (**Supplementary Note 4**), which contribute to the extraction of sharp edge and along-edge continuity feature in the third dimension, respectively. **Extended Data Fig. 1a** and **b** show the principle and workflow of the MRA deconvolution algorithm, which can be applied to all kinds of fluorescence imaging techniques as a post-processing method.

### High-fidelity of MRA deconvolution

Although most deconvolution methods can improve the resolution and provide a visually appealing image, it is of great importance to maintain fidelity in fluorescence microscopy: the result should be close to the ground-truth (GT), and the derived sharp image should be able to revert to the blurry outcome by re-convolving with the actual point spread function (PSF). To evaluate the fidelity, we synthesized some simple geometrics as GT images, which were then blurred and added with Gaussian noise to test the effectiveness of the MRA deconvolution algorithm (**Supplementary Note 5**, **Supplementary Figs. 4-6**). The results show that the MRA deconvolution algorithm can faithfully recover these geometrical structures polluted by severe noise. By tuning the illumination power from 3% to 100%, we captured raw SIM images of U2OS cell actin filaments with varying SNR (**Fig. 1b**). The 2D-MRA deconvolution resolves the dense actin network from noise contamination and is robust to the noise level. The resolution would be greatly damaged with the improvement of SNR in conventional deconvolution algorithms (**Supplementary Fig. 1**). Owing to the superior signal-noise distinguishing ability of framelet and curvelet, the 2D-MRA deconvolution improves the SNR of the actin image with little contamination in resolution. Two parallelly aligned actin filaments can be resolved by the MRA deconvolution even with the 3% power illumination, whereas conventional SIM reconstruction can barely resolve it until the illumination power reaches 20% (**Fig. 1c**). Using only framelet, curvelet, or traditional orthogonal wavelet sparsity as regularization cannot reach the same image quality as MRA (**Supplementary Fig. 7**).

**Fig. 1.**
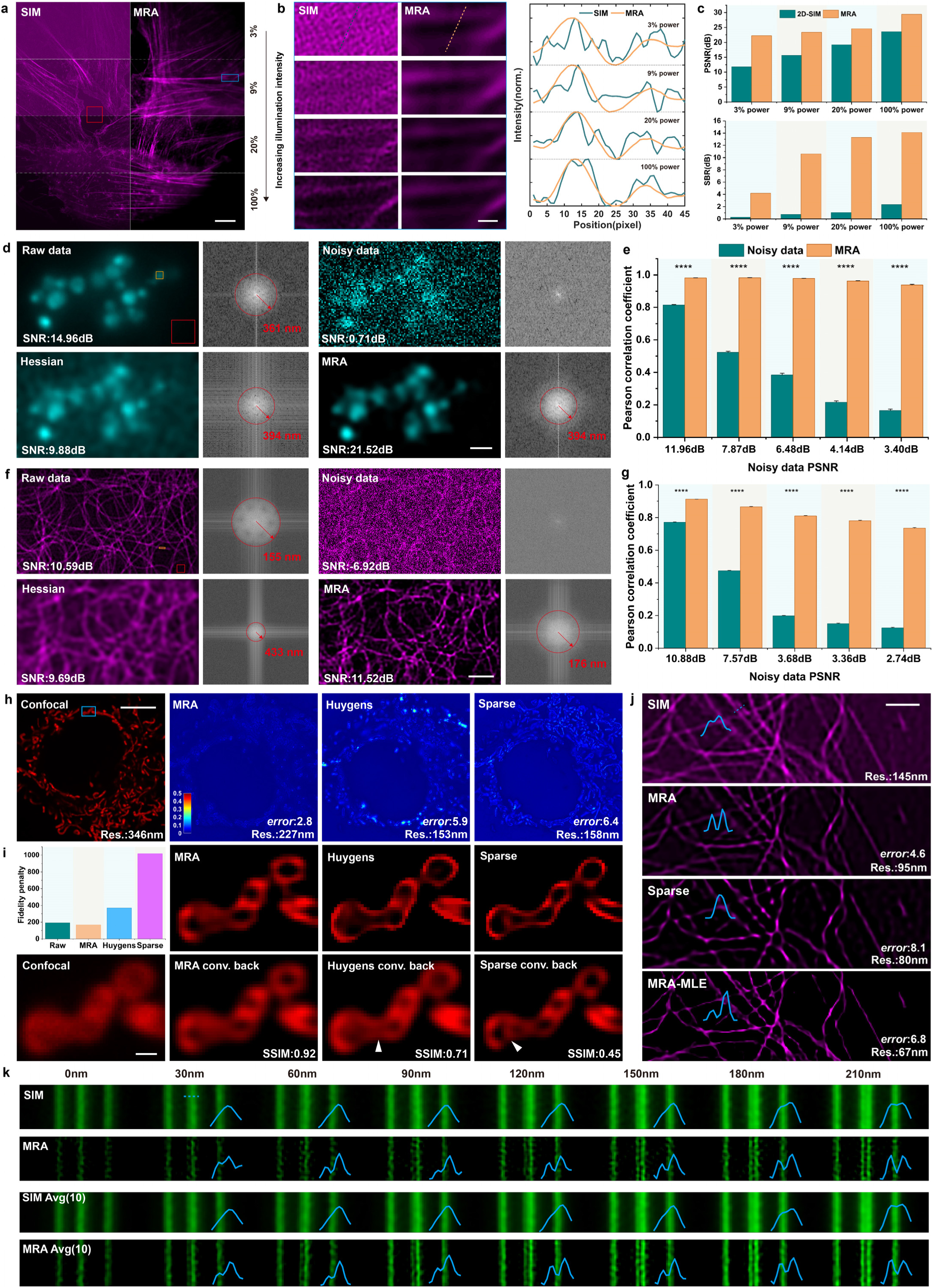
MRA deconvolution recovers information from ultralow-SNR fluorescence images and supports physical resolution improvement. **a.**The SIM images of actin in U2OS cell under different illumination intensities are shown in the left column, and the corresponding MRA deconvolution results are shown in the right. **b.** The magnified blue boxed region in b (Left), and the intensity profile plots of raw SIM images and MRA deconvolved images along the lines indicated by the left column dotted lines (Right). **c.** The PSNR of the original SIM and MRA deconvolved images using the red-boxed region in b as the noise and background sample (Top), and the SBR of the blue boxed region in a (Bottom). **d, f.** The raw data are the middle frame of the lysosome and microtubule time-lapse images (*n*=20). Noisy data are raw images contaminated with Gaussian additive noise. The corresponding MRA and Hessian deconvolution results are shown at the bottom. The signal is estimated through the orange boxed region, and the noise is estimated through the red boxed region, which is used to calculate SNR. **e, g.** Pearson correlation between MRA deconvolved data and raw data under multiple Gaussian noise levels (*n*=20). ****, *p*<0.0001. **h.** The confocal image of mitochondria in COS-7 cell mitochondria, and the resolution-scaled error map of the MRA, Huygens, and Sparse deconvolution result obtained by NanoJ-SQUIRREL software. **i.** Fidelity penalty of each image, and magnified blue boxed regions in h. The PSF for deconvolution is used to convolve the deconvolved image back (marked by conv. back). **j.** SIM image of microtubules in COS-7 cell and its deconvolution results by MRA, Sparse, and MRA-MLE. **k.** Separation of Argo-SIM parallelly aligned fluorescence lines by MRA deconvolution. The bottom Avg (10) denotes averaging ten raw images to improve SNR and then processing with MRA deconvolution. The resolution displayed in h, j is estimated through decorrelation analysis^45^, and in d, f is estimated by the spectrum edge due to noise influence on the decorrelation analysis. Scale bars, 10 *μ*m (h), 5 *μ*m (a), 2 *μ*m (f), 1 *μ*m (d, g, j), 0.5 *μ*m (b, i).

To assess the performance of MRA on continuous frames, we captured the temporal fluorescence image sequence of lysosomes and microtubules as a reference, which was then contaminated with Gaussian additive noise. We then deconvolved the noisy image sequence using Hessian and MRA deconvolution, respectively. Considering the diversity of parameter selection, we compare the denoising performance of the two algorithms under equivalent SNR or resolution level. When deconvolving the lysosome image with equivalent resolution (**Fig. 1d**), the SNR of the MRA deconvolved image is >10 dB higher than Hessian deconvolved image, indicating the superiority of MRA regularization in signal-noise discrimination. When the SNR of the Hessian deconvolved microtubule image is still >1 dB lower than the MRA deconvolved result, the resolution of the Hessian deconvolved image, as estimated by the Fourier spectrum edge (**Methods**), deteriorates to ∼1/3 of the original image (**Fig. 1f**). The Hessian deconvolved image exhibited obvious signal blurring, which was not observed in the MRA deconvolved image, where the important edge information was preserved in the deconvolution process. We further test the performance of MRA deconvolution with multiple Gaussian noise levels (**Fig. 1e**, **g**, **Extended Data Fig. 2a**, **b** and **Supplementary Videos 1**, **2**), where we increase the value of regularization parameters with the increase of noise level to improve the denoising ability. The results show that compared with the noisy data, the images deconvolved by MRA have significantly higher Pearson correlation with the raw data, and are robust to various noise levels. Next, we test the resolution-maintaining ability of MRA by holding the regularization parameter value (**Extended Data Fig. 2c**, **d**), which generates images with equivalent resolution but lower SNR for higher noise level. Setting the ultra-strong correlation (Pearson correlation >0.8) between raw data and MRA deconvolved data as the criteria, MRA deconvolution can recover the lysosome and tubulin structure with an unhindered resolution, even when the image SNR is as low as ∼ -2 dB and -10 dB, respectively.

We are curious about whether MRA regularization can support physical resolution improvement (i.e., directly minimizing the fidelity term ||*f*-*Ax*||2), which is generally suboptimal for real microscopic images due to its fragility of noise and incompatibility with variance-based noise-control regularization since the high-resolution image is always non-smooth. To test this, we used confocal microscopy to capture a COS-7 cell mitochondria image, which shows a >1.5-fold physical resolution improvement after MRA deconvolution with a relatively low regularization parameter value. We then compare the performance of this physical deconvolution method with the advanced statistical MLE method using NanoJ-SQUIRREL error analysis^46^. The results show that though Sparse and commercial Huygens deconvolution provides numerically higher resolution, their resolution-scaled error is several times higher than MRA. Convolving the deconvolution result back, artificial structures appear in MLE convolved-back image while not in MRA (**Fig. 1o**). We calculated the fidelity penalty ||*f*-*Ax*||2 of each resolution-enhanced image, and found that only MRA reduces the penalty value (**Fig. 1o**), which reveals the difference between the physical and statistical method. We further conducted a quantitative analysis of the two deconvolution methods on a high-SNR SIM microtubules image (**Extended Data Fig. 3**), which shows that MLE methods over-inferring the full-width-at-half-maxima (FWHM) of tubulin filament by >60 nm. In contrast, although visually less appealing, MRA correctly provides ∼40 nm FWHM reduction to the tubulin filament with respect to the PSF, as confirmed by its convolved back image. We also evaluated the effect of iteration times on the deconvolved FWHM value (**Extended Data Fig. 3d, e**), which shows that as MRA converges to the theoretically unblurred FWHM value with increasing iteration times, whereas MLE iteration arbitrarily reduces the FWHM value even many times lower than the theoretical limit.

We also investigate the impact of regularization on the two deconvolution methods by fabricating MRA-MLE that uses MRA denoising to support the RL iteration (**Supplementary Note 3.3**). On synthesized geometrics images, the lines and rings can be resolved by MRA-MLE deconvolution with higher numerical resolution than MRA, but all are severely twisted with obvious artifacts even with relatively low noise levels (**Supplementary Note 5** and **Supplementary Figs. 8-10**). We speculate that both the artifact-proneness feature and the distance between the denoised and real image may influence the performance of the MLE method. Benefiting from MRA regularization, MRA-MLE outperforms Sparse deconvolution on SIM microtubule and actin images with reduced artifacts and better-preserved details even at a higher resolution level (**Fig. 1n** and **Supplementary Fig. 11**). However, the error of MRA-MLE image is still higher than MRA approach with apparent discontinuity and abnormal edges. Moreover, the impact of parameters is considerable in the denoise-MLE scheme: the denoising parameter influences the information transmitted to MLE iteration, and different iteration times yield rather different results. Conversely, the parameter tuning in MRA only regulates the image SNR and resolution without affecting the physically extracted high-frequency information. Therefore, we believe that physical deconvolution is optimal for fluorescence image processing since high fidelity and objectivity are rather crucial. In this work, we mainly explore the utilization of MRA. To clarify the maximum resolving ability of MRA, we image the Argo-SIM resolution test panel under 1.49-NA SIM with good SNR (∼15 dB), allowing a low regularization value to strengthen the resolution improvement. Surprisingly, MRA can readily resolve the 30 nm-distance parallel line (**Fig. 1k**). By averaging ten images to enhance image SNR (∼17 dB), MRA offers better signal quality.

### MRA deconvolution assists live-cell imaging

With better prior knowledge, the MRA deconvolution algorithm demonstrates excellent performance in recovering information from photon-limited fluorescence images. Endoplasmic reticulum (ER)^47,48^ is a large organelle that plays a crucial role in a variety of life activities such as the synthesis and modification of proteins. To capture the high dynamics of ER network, a high temporal resolution is often required. MRA can help improve the imaging speed by compensating for the signal loss due to short exposure time. We applied MRA deconvolution algorithm to assist the rapid wide-field imaging of ER tubules labeled by KEDL-GFP plasmid (**Fig. 2a** and **Supplementary Video 3**) with an exposure time of ∼ 2.4 ms, corresponding to a temporal resolvability of ∼ 417 Hz. The illumination intensity was only ∼2.38 W/cm^2^, resulting in a low SNR of ∼1 dB. Due to the ultralow SNR, the ER network in the original image cannot be correctly segmented even by some advanced machine-learning methods^49^. After MRA deconvolution, the ER network can be finely segmented (**Fig. 2a**). The information submerged in severe noise has been successfully resolved by MRA deconvolution, through which the rapid movement of vesicles and variation of ER network can be finely observed (**Fig. 2b**).

**Fig. 2.**
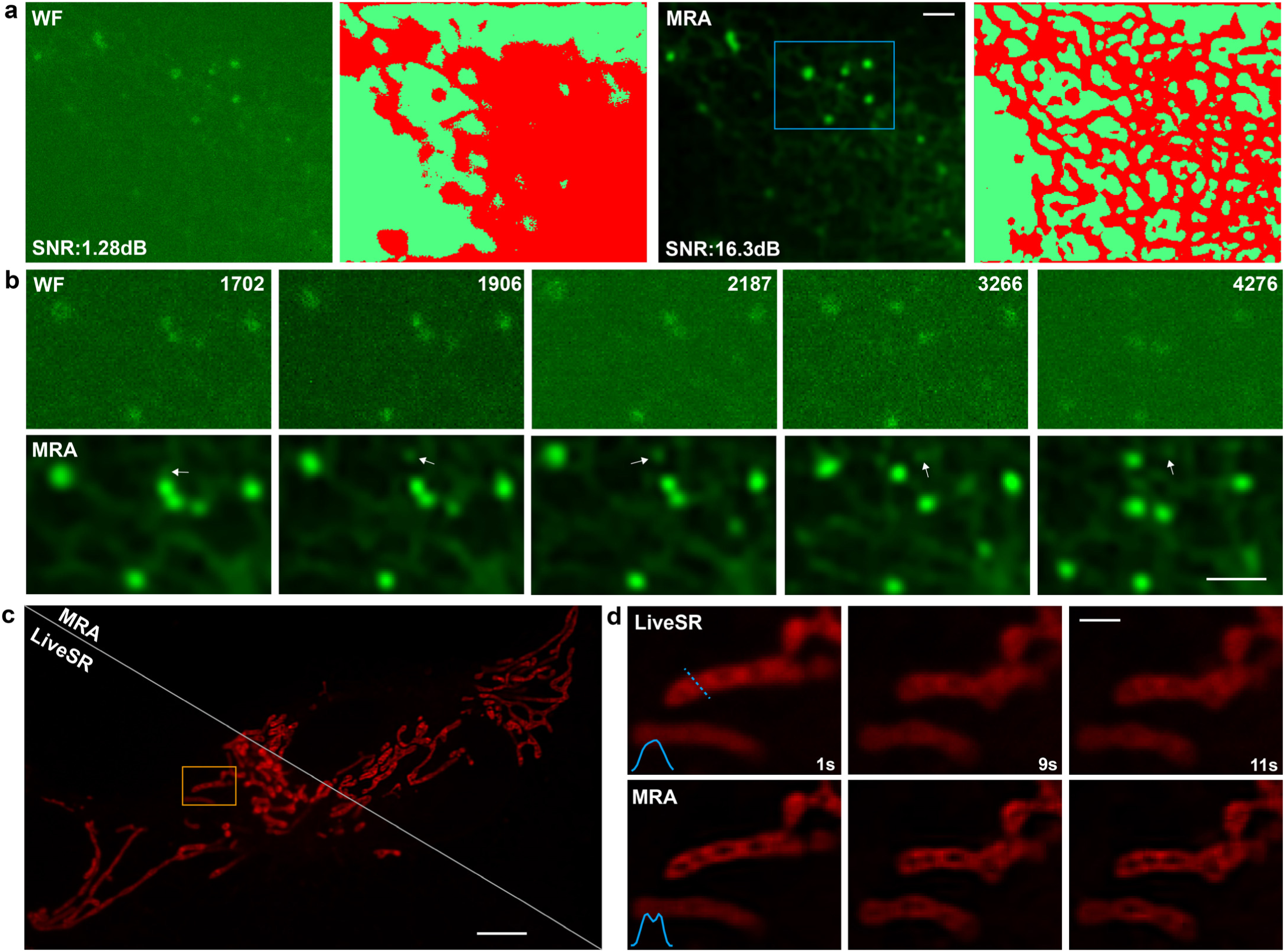
MRA deconvolution assists live-cell imaging. **a.** Wide-field time-lapse images of ER tubules in COS-7 cell labeled by KEDL-GFP plasmid, and the MRA deconvolution result. The imaging contains a total frame of 4,500 with a frame rate of ∼ 417 fps. The ER network is segmented using TWS^49^, a machine-learning toolbox for organelle segmentation. **b.** The magnified blue boxed region in a. The number on the right top denotes the frame number. **c.** LiveSR time-lapse images of mitochondria in U2OS cell, and the MRA deconvolution result. **d.** The magnified orange boxed region in c. Scale bars, 5 *μ*m (c), 2 *μ*m (a, b), 1 *μ*m (d).

Mitochondria are highly dynamic organelles in mammalian cells that play a crucial role in energy support of cell lifecycle^50,51^. To investigate mitochondrial activities, a fine balance between resolution and toxicity is needed. Some subtle structures of mitochondria, such as the cristae, can be resolved by SR STED imaging technique, which on the other hand induces severe toxicity that hampers the mitochondria viability. While even for some SR techniques with lower toxicity, such as commercial LiveSR imaging, it is still difficult to clearly capture the cristae dynamics for insufficient resolution (**Fig. 2c**, **d**). The MRA deconvolution can help improve the resolution of LiveSR mitochondria image with high-fidelity and resolve the cristae structure submerged by optical blurring (**Fig. 2e**). Benefited from the high-fidelity feature of MRA deconvolution, there is no artifact introduced in the resolution-improved image. Tuning parameters in MRA only regulates the image SNR and resolution, which does not influence the inferred high-frequency information. In comparison, the MLE-based method shows rather different results as the change of denoising parameters and MLE iteration times (**Supplementary Fig. 12**).

### DeepMRA deconvolution algorithm

Although MRA deconvolution achieves satisfactory results in resolving biological structures from severe noise contamination and enhancing the image resolution with high fidelity, it is inherently limited to dealing with fluorescence background. Traditional ||*x*||1 regularization^10,52^ indiscriminately attenuates the entire image, which can enhance the image contrast and remove uniform background. However, if the background intensity exceeds the signal, this method will instead enhance the background. Although some preliminary background subtraction methods can alleviate this issue, these operations inevitably reduce image SNR, resulting in the loss of high-frequency or low-intensity details in the subsequent deconvolution process (**Extended Data Fig. 4**). Furthermore, when denoising low-SNR images, traditional deconvolution algorithms can amplify the background and uneven intensity distribution, exacerbating the problem. To address this issue, here we provide a novel scheme to solve this problem by introducing a bias thresholding mechanism to the MRA deconvolution iteration, termed DeepMRA deconvolution:

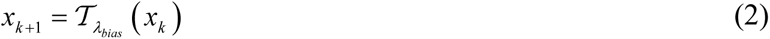

where *x_k_* denotes the image in the *k*th iteration, *T* denotes the soft-thresholding operator, *λ_bias_* is the bias thresholding matrix that penalizes the background component. The bias thresholding matrix *λ_bias_* is generated based on the weighted addition of the framelet-coarsened and the original image, which simultaneously reflects the low-frequency background and the fluctuation of the original image. The complete procedure for fabricating bias thresholding matrix *λ_bias_* is summarized in **Supplementary Note 3.2**. This method selectively penalizes the background information and controls the noise in each deconvolution iteration, enabling background mitigation with preserved details. Because in the deconvolution iteration, the trade-off between background attenuation and MRA contribution is still inherent, we add a preliminary background subtraction step before the spatiotemporal continuity denoising step to alleviate ultra-strong fluorescence background (**Supplementary Note 3.2** and **Supplementary Fig. 13**). The whole procedure of DeepMRA deconvolution is shown in **Extended Data Fig. 1c**.

With its remarkable background inhibition ability, DeepMRA deconvolution has substantially expanded the application scope of MRA deconvolution. We demonstrate the feasibility of DeepMRA deconvolution in various fluorescence microscopies and samples. We acquired images of mouse kidney cell nuclei using both wide-field and SD-confocal microscopy, which exhibited significant background interference (**Fig. 3a**). Upon deconvolution with DeepMRA, the images showed a substantial reduction in background with finely preserved details. The DeepMRA deconvolved wide-field image even yields better visual perception than the original SD-confocal image. We then imaged the actin structure of mouse kidney cells using Nikon SoRa mode, which was noticeably affected by out-of-focus signals. After DeepMRA deconvolution, the noise and background were significantly reduced, leading to better 3D rendering results (**Fig. 3c** and **Supplementary Video 4**). Background interference can significantly limit MRA deconvolution, especially when there is strong noise and a large regularization parameter value is required, hindering the extraction of high-frequency information (**Fig. 3c**). With the bias thresholding scheme, DeepMRA can overcome this barrier and extract rich details from fluorescence images contaminated by background and noise. This feature allows DeepMRA to provide 1.6-fold physical resolution improvement to the SD-confocal mitochondria image, and 2.2-fold to the SoRa image contaminated by background (**Fig. 3d**). The DeepMRA deconvolved SD-confocal image exhibits high similarity with the SoRa image, while artifacts appear in the MLE deconvolved image even at equivalent resolution level (**Supplementary Fig. 14**). We further verify the fidelity of DeepMRA resolution enhancement in other low-high resolution imaging pairs with background (**Extended Data Fig. 5**). Additionally, we demonstrate the effectiveness of DeepMRA deconvolution for organelles and cytoskeletons in other common fluorescence setups, including LiveSR (**Fig. 3e**), SIM (**Fig. 3f**), STED (**Fig. 3g**), and light-sheet microscopy (**Extended Data Fig. 8**).

**Fig. 3.**
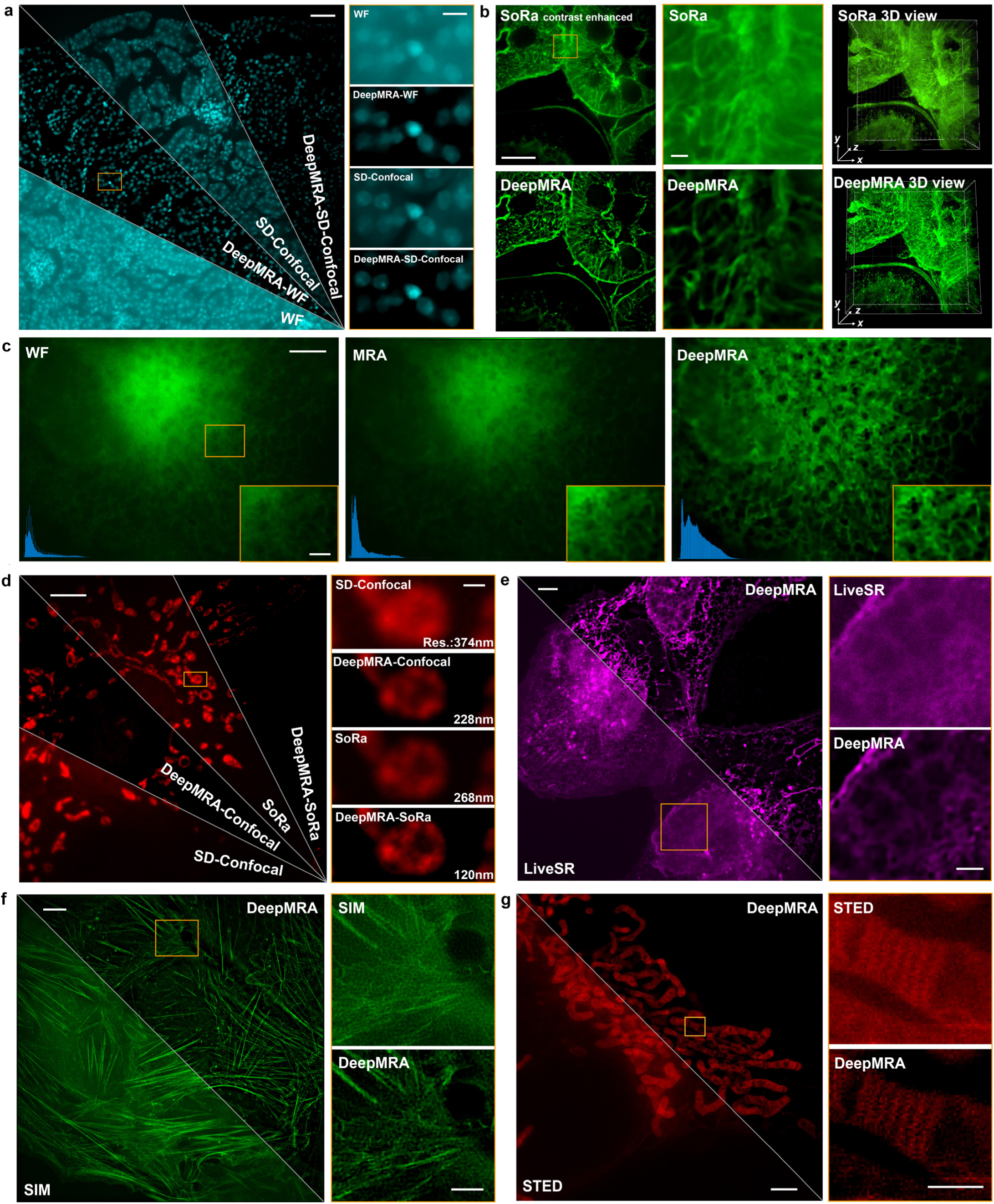
DeepMRA enhances fluorescence image with background in various microscopies. **a.** Wide-field and SD-confocal image of nuclei in mouse kidney cell, and the corresponding DeepMRA deconvolution results (Left). Magnified orange boxed region in the left column (Right). **b.** SoRa image of actin in mouse kidney cell, and the DeepMRA deconvolution results (Left). The magnified orange boxed region is in the left column (Middle). The 3D view of the SoRa and DeepMRA deconvolution result (Right). **c.** ER tubules in COS-7 cell captured by wide-field microscopy, and the MRA and DeepMRA deconvolution results, the blue graph in the left bottom shows the histogram. The magnified orange boxed region is shown at the right bottom. **d.** BPAEC cell mitochondria image captured by SD-confocal microscopy and SoRa, and the corresponding DeepMRA deconvolution results (Left). Magnified orange boxed region in the left column (Right). **e.** The LiveSR captured Nile-Red labeled U2OS cell image, and the DeepMRA deconvolution result (Left). Magnified orange boxed region in the left column (Right). **f.** The SIM image of actin in U2OS cell, and the corresponding DeepMRA deconvolution result. Magnified orange boxed region in the left column (Right). **g.** The STED image of mitochondria in COS-7 cell, and the DeepMRA deconvolution result (Left). Magnified orange boxed region in the left column (Right). Scale bars: 50 *μ*m (a Left), 10 *μ*m (a Right, b Left), 5 *μ*m (c, d Left, e Left, f Left, g Left), 2 *μ*m (c subgraph, e Right, f Left, g Left), 1 *μ*m (b Right), 0.5 *μ*m (d Right, g Right).

### Long-term organelle interaction imaging assisted by DeepMRA

As a general algorithm for enhancing fluorescence images, DeepMRA has great potential to support research in a variety of life science fields. Here we demonstrate the application of DeepMRA deconvolution to long-term fluorescence imaging of organelle interactions, which is a critical aspect of cellular function^48,53^. We first performed two-color wide-field imaging of microtubules and lysosomes in COS-7 cell labeled by LysoView^TM^ 488 and SiR-Tubulin Kit (**Fig. 4a** and **Supplementary Video 5**). After applying DeepMRA deconvolution with appropriate parameters, background and noise are effectively removed and the resolution is improved by ∼ 86 nm (**Fig. 4b**), enabling better observation of the contact between tubulins and lysosomes (**Fig. 4c**). We quantified organelle colocalization by calculating Mander’s overlap and Pearson correlation coefficient, both of which show stable bonding during the ∼12-minute recording period (**Supplementary Fig. 15**). With the enhanced image, we were able to successfully segment the structures of tubulins and lysosomes and determine their contact region (**Fig. 4d**), a task that was difficult with the raw image due to background and noise interference. Additionally, we performed rapid wide-field imaging of co-labeled microtubules and ER tubules, where DeepMRA effectively recovered lost information caused by low photon dose and facilitated visualization of the contact region between the two organelles (**Extended Data Fig. 7 and Supplementary Video 6**).

**Fig. 4.**
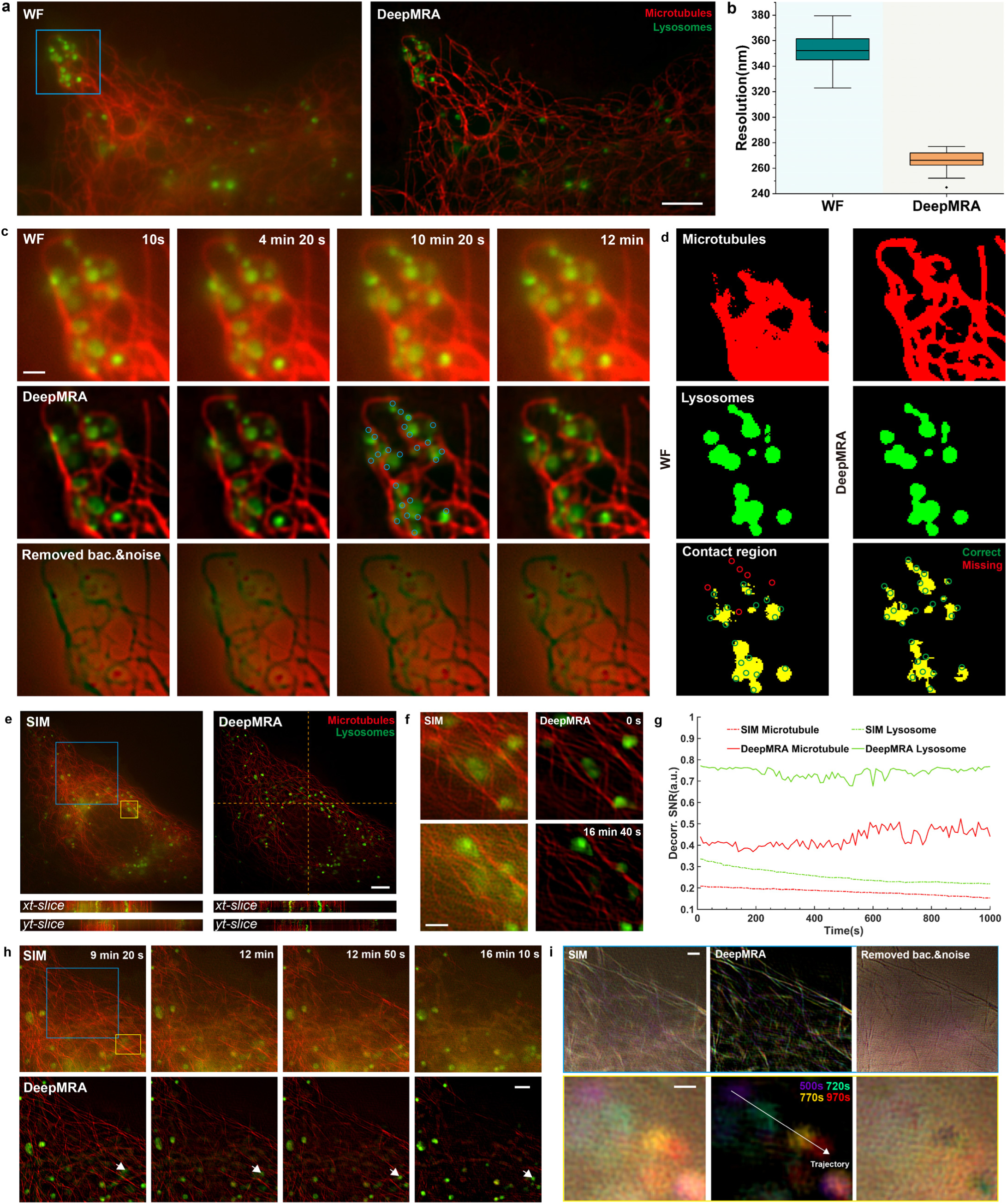
DeepMRA deconvolution assists long-term fluorescence imaging of microtubule-lysosome interaction. **a.** Dual-color wide-field time-lapse images of microtubules (red) and lysosomes (green) in COS-7 cell, and the DeepMRA deconvolution result. The imaging lasts 12 minutes and 20 seconds with 74 frames. **b.** Resolution of the original wide-field and DeepMRA deconvolved microtubule image stack. **c.** Magnified blue boxed region in a. **d.** Segmented microtubules and lysosomes image at 10 min 2 s time point through direct thresholding. The thresholding value for normalized wide-field tubulin and lysosome images are 0.33 and 0.38, and for MRA are 0.11 and 0.2. The contact region is determined by the multiplication of the two segmented images. The green and red circles indicate the correctly determined and missing contact sites based on the manual mark in c. **e.** Dual-color SIM time-lapse images of microtubules (red) and lysosomes (green) in COS-7 cell, and the DeepMRA deconvolution result. The imaging lasts 16 minutes and 40 seconds with 100 frames. **f.** Magnified yellow boxed region in a. at 0 and 1,000 second time point. **g.** The SNR at different time points is estimated by decorrelation analysis. **h.** Magnified blue boxed region in a. **i.** Temporal projection of the blue boxed region in d with the microtubule channel (Top), and the yellow boxed region in d with the lysosome channel (Bottom). Scale bars: 5 *μ*m (a, e), 2 *μ*m (h), 1 *μ*m (c, f, i Top), 0.5 *μ*m (i Bottom).

In dual-color imaging of microtubules and lysosomes using SIM, we also demonstrate the efficacy of DeepMRA (**Fig. 4e** and **Supplementary Video 7**). Through its deconvolution process, DeepMRA effectively reduces the noise and recovers signal loss due to photobleaching (**Fig. 4f**, **g**). We were able to observe the depolymerization process of microtubules, which is presumably due to mitosis (**Fig. 4h**). During this process, some lysosomes gradually shifted from the microtubule accumulation region. However, the raw SIM image suffered from excessive background and noise, which hinders our observation. Fortunately, DeepMRA was able to precisely eliminate this interference while preserving the original signal, thus enhancing our ability to visualize the dynamic movements of tubulin and lysosomes over time (**Fig. 4i**). Notably, conventional deconvolution algorithms typically sacrifice low-intensity and high-frequency details in the process of enhancing the densely meshed microtubule image (**Extended Data Fig. 4a**).

Subsequently, we employed a commercial LiveSR microscope to image the interaction between mitochondria and microtubules. Due to the sensitivity of both organelles to illumination, we utilized low-intensity illumination to ensure their viability (**Fig. 5a** and **Supplementary Video 8**). Nevertheless, the tubulin signal was plagued by noise and the presence of undesired ultrabright regions hindered accurate observation. With the assistance of DeepMRA, we were able to recover the tubulin signal and alleviate uneven intensity distribution by using an intentionally selected bias parameter, enabling us to observe the mitochondrial fission process around the microtubule (**Fig. 5b**). In contrast, conventional algorithms typically amplify the undesired ultrabright regions (**Extended Data Fig. 4c**). Our method also enabled us to visualize two consecutive mitochondrial fission and fusion events around the microtubules in the LiveSR system (**Extended Data Fig. 8** and **Supplementary Video 9**).

**Fig. 5.**
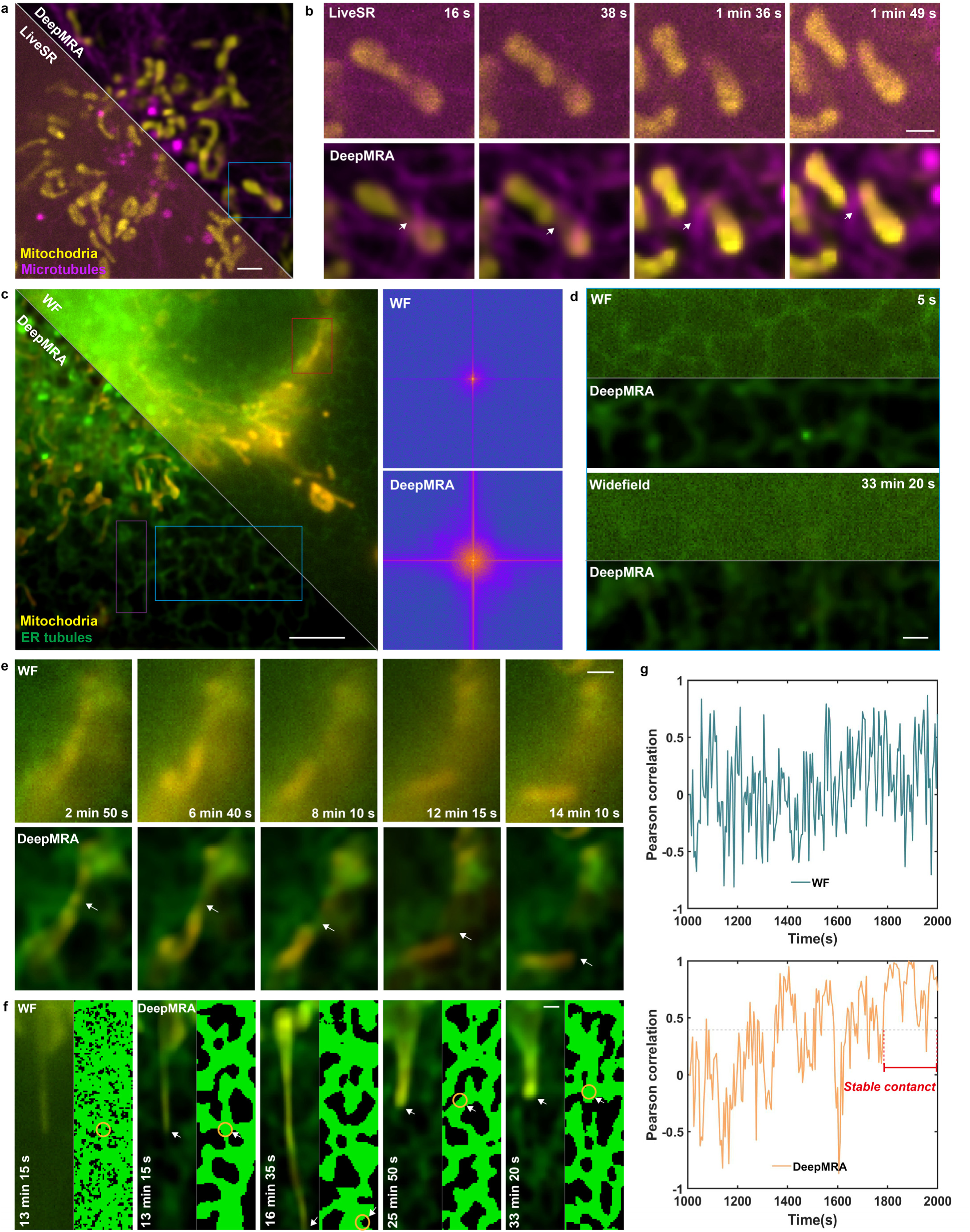
DeepMRA deconvolution assists long-term fluorescence imaging of mitochondria interaction with microtubules and ER tubules. **a.** Dual-color LiveSR time-lapse images of mitochondria (yellow) and microtubules (magenta) in U2OS cell, and the corresponding DeepMRA deconvolution result. The imaging lasts 2 minutes with 120 frames. **b.** Magnified blue boxed region in a at different time points. The white arrow denotes the mitochondrion fission site. **c.** Dual-color wide-field time-lapse images of mitochondria (yellow) and ER tubules (green) in COS-7 cell and the DeepMRA deconvolution result (Left). The imaging lasts 33 minutes and 20 seconds with 400 frames. The Fourier spectrum of wide-field and DeepMRA deconvolved images (Right). **d.** Magnified blue boxed region in g at 5 s and 33 min 20 s time point (Right). **e.** Magnified red boxed region in c, which shows a mitochondrial fission process. **f.** Magnified purple boxed region in c. The right column in each graph shows the segmented ER network by the TWS method and the position of the mitochondrial distal end. **g.** Pearson correlation between the ER tubules and mitochondrial distal end at different time points. Scale bars: 5 *μ*m (c), 2 *μ*m (a, d), 1 *μ*m (b, e, f).

Fluorescence imaging in the vicinity of the nucleus region can be severely affected by out-of-focus signal interference due to the greater thickness and dense cellular structure. To investigate organelle interactions in this region, microscopy systems with better optical sectioning capabilities are usually required as severe fluorescence background often exists. However, these imaging systems entail additional costs and may compromise on toxicity and imaging speed. Here, we demonstrate that by utilizing the computational sectioning and denoising abilities of DeepMRA, interactions between mitochondria and the ER tubules near the nucleus region can be observed with great precision even using the simplest wide-field microscopy. We employed a low illumination intensity (∼2.33 W/cm^2^ for the ER channel and ∼3.26 W/cm^2^ for the mitochondria channel) to support imaging for more than 30 minutes without severe photobleaching and phototoxicity effects (**Fig. 5c,d** and **Supplementary Video 10**). Disrupted by excessive noise and background information, the ER signal in the original image is almost invisible, which can be finely resolved by DeepMRA while conventional algorithms are incompetent (**Extended Data Fig. 4b**). With the assistance of DeepMRA, we observed a mitochondrion division event near the ER network (**Fig. 5e**), which has also been observed by many SR imaging techniques^48^. We also observed a mitochondrion extension process, where the mitochondrion extended to a length of ∼ 5.2 *μ*m, then gradually shrunk and attached to a surrounding ER network (**Fig. 5f**). We quantitatively analyzed their interaction by calculating the Pearson correlation between the mitochondrial distal end and the ER tubules (**Fig. 5g**). During the extension and shrinkage process, peaks and valleys of the Pearson correlation curve were formed as the mitochondrion shuttled through the ER network. Eventually, it was observed that the mitochondrion distal end and the ER network showed stable contact (**Supplementary Video 10**). While the Pearson correlation calculated through the raw wide-field images randomly varied during this process due to noise and background interference, the DeepMRA-enhanced image revealed the close bond of the two organelles in the last frames with a continuously high Pearson correlation (>0.4).

## Discussion

The desire to improve the spatiotemporal resolution and reduce the toxicity of fluorescence microscopy is long-lasting due to the inherent photon limit of fluorophores. The computational approach has been an increasingly important driving force to meet such expectations. Ideally, there is no limit in resolution in noiseless imaging because we can always get the unblurred image through direct inverse filtering. However, in practical fluorescence imaging the noise is abundant, and eliminating the noise relies on our assumption about the fluorescence image feature. In this work, we abandon the conventional scheme that characterizes the continuity feature of the biological images using naïve spatial or Fourier domain analysis. Instead, we characterize the sharp-edge contrast and along-edge continuity feature of fluorescence image by regularizing its sparsity in the framelet and curvelet domain, respectively. We apply a similar idea to denoise temporal sequence and 3D imaging, in which the sharp-edge contrast and along-edge continuity feature are subtracted by regularizing the 3D framelet and DTCW domain sparsity, respectively. Our method, termed MRA deconvolution, shows superior performance than conventional variance-based regularization because we introduce the prior knowledge to strengthen edge information and distinguish the continuity of the biological itself and its surrounding background, which allows noise removal with finely preserved details. Furthermore, the superior MRA regularization supports physical resolution improvement under noise condition, which is highly appreciated in fluorescence image processing since high fidelity is required. In various fluorescence imaging systems, we demonstrate utilization of MRA deconvolution, which helps separate Argo-SIM 30-nm parallelly aligned lines, recover signals from ultra-low SNR ER image due to short exposure, and resolve mitochondrial cristae in moderate SR imaging system.

Despite the excellency in denoising and resolution enhancement, MRA deconvolution still encounters difficulty when dealing with images with strong background. Introducing preliminary background subtraction or ||*x*||_1_ regularization to deconvolution may help enhance contrast and mitigate background, which, however, would lose high-frequency or low-intensity details and lose efficacy when the background is ultra-strong. This motivates the development of DeepMRA deconvolution, which estimates background using the MRA approach and eccentrically attenuates the background in each iteration. This novel deconvolution canonical form has a better performance because the background is iteratively subtracted rather than once unprecise estimation, and the noise is controlled in each iteration. Moreover, by tuning the parameter, DeepMRA can also be utilized to correct uneven intensity. With this feature, DeepMRA can enhance fluorescence images in various microscopies and imaging conditions. We examine the performance of DeepMRA in multiple fluorescence imaging setups, including wide-field, confocal, SD-confocal, light-sheet/super-resolution LiveSR, SoRa, SIM, and STED, in which DeepMRA effectively mitigates background, removes noise and enhances resolution while finely preserving original details. Besides the biological cell, DeepMRA is also effective when imaging thick sample, such as the mouse kidney slice demonstrated in this work.

We also provide the MATLAB source code for developers and also develop interactive software of the two deconvolution algorithms for users. The MRA deconvolution is a standard fluorescence image recovery algorithm, while DeepMRA is bestowed with some enhancement nature. We hope that as an extension of the deconvolution algorithm family (**Supplementary Fig. 16**), MRA and DeepMRA with noise-robustness and the high-fidelity feature can help assist life science research. To help master the utilization of MRA and DeepMRA, we provide a detailed parameter tuning guidance (**Supplementary Note 6** and **Supplementary Figs. 17-21**) and demonstrate the effect of each parameter (**Supplementary Figs. 22-25**). The parameter tuning essentially is the trade-off of image resolution and SNR, while the inferred high-frequency information is irrelevant to parameter choice in MRA, making it a stable and reliable tool for enhancing life science images. Moreover, we are also optimistic about the extension of the two algorithms. The astonishing resolution enhancement and background inhibition of wide-field microscopy may be beneficial for some event-trigger microscopy setups^55,56^ as they employ wide-field imaging to monitor cell activities and determine the activation of high-resolution SR imaging. The combination of MRA with the deep-learning model is also worthy of expectation. MRA can provide physically enhanced high-SNR images for model training, and improve the resolution after deep-learning denoising. Introducing MRA-based prior knowledge to the deep-learning model may improve the performance, as some recent works indicate the effectiveness of incorporating some analytical models to the deep-learning network^14,16^.

## Methods

### Fluorescence microscopes

We employed various fluorescence microscopes to test the effectiveness of the MRA and DeepMRA deconvolution algorithms. The commercial Airy Polarization-SIM super-resolution microscopy system (Airy, China) is used to capture the wide-field and SIM images displayed in Fig. 1,Fig. 2,Fig. 3c,f,Fig. 4,Fig. 5, Extended Data Fig. 3,Extended Data Fig. 5 and Extended Data Fig. 6. The Zeiss confocal laser scanning microscopy LSM 980 with Airyscan 2 (Zeiss, Germany) is used for imaging the mitochondria displayed in Fig. 1i. The Nikon CSU-W1 SoRa spanning-disk microscopy system (Nikon, Japan) is used to capture the wide-field,SD-confocal, and SoRa image displayed in Fig. 3 a, b and d. Yokogawa spinning disk equipped with a LiveSR super resolution module (Gataca systems, France) is employed to capture the LiveSR image displayed in Fig. 2,Fig. 3,Fig. 5,Extended Data Fig. 7. Leica SP8 STED 3X microscope (Leica, Germany) is employed to capture the STED image displayed in Fig. 3. A detailed imaging parameter of each image is listed in Supplementary Table 1.

### Cell maintenance and preparation

COS-7 cells and U2OS cells were cultured high glucose medium DMEM (Gibco, 11995-040) with the addition of 10% fetal bovine serum (FBS, Gibco, 10099) and 1% penicillin-streptomycin antibiotics (10,000U/mL, Gibco, 15140148), in an incubator at 37 °C with 5% CO^2^. For live cell imaging experiments, cells were seeded in μ-Slide 8 Well (ibidi, 80827). For fixed-cell imaging experiments, cells were seeded on coverslips (Thorlabs, CG15CH2). The imaging sample was prepared until the cells reached a confluency of 75%.

### Fixed sample

#### Mouse kidney section sample

The specimens of phallodin-AF568-labeled actin in mouse kidney section were commercially available (FluoCells Prepared Slide #3, Invitrogen, F24630).

#### BPAEC cell sample

We use commercial FluoCells ™ Slide #1 (ThermoFisher, F36924) to test the performance of our algorithms. It contains BPAEC cells stained with MitoTracker ™ Red CMXRos, Alexa Fluor ™ 488 ghost pen cyclic peptide and DAPI.

#### Argo-SIM standard slide

In order to verify the resolution and fidelity of the algorithm, commercial fluorescent standard samples (Argo-POWER SIM Slide V2, Argolight) were photographed using Airy-SIM. The ground truth image consists of pairs of fluorescent doublets (spacing from 0 nm to 390 nm; *λ*ex = 488 nm).

#### Labeling actin in fixed U2OS cells

The cell was fixed with 4% formaldehyde (R37814, Invitrogen) for 15 minutes at room temperature. After washing the sample with PBS, we permeabilize samples with 0.1% Triton™ X-100 (Invitrogen, HFH10) for 15 minutes. After washing with PBS, we used Alexa Fluor™ 568 Phalloidin (Invitrogen, A12380) / Alexa Fluor™ 488 Phalloidin (Invitrogen, A12379) dye to stain the actin filament for 1h at room temperature. Place coverslips in a covered container to prevent evaporation during incubation. Wash the samples two or more times with PBS.and place the stained coverslips in a dark place to dry naturally. The coverslip was sealed with 30 µL of Prolong (Invitrogen, P36984) mounting medium and placed at 4°C overnight to air-dry and then observe.

#### Labeling membrane structures in fixed U2OS cell

To label all lipid membrane structures in the cell, 1 ug/ml Nile Red (Invitrogen, N1142) was added into the culture medium 1 h before imaging and was present during imaging.

#### Centrosome sample for expansion microscopy

The Expansion microscopic sample of the centrosome was a gift from Fu’s laboratory. The associated sample preparation procedure and imaging method have been described previously^57^.

### Live-cell sample

#### Transfection of GFP-KDEL plasmid to mark endoplasmic reticulum

To label the dynamic structure of the ER in living cells, we transfected GFP-KDEL to COS-7 cells with Lipofectamine 3000 (Invitrogen, L3000) transfection reagent. The imaging was performed 36-48 hours after transfection.

#### Labeling mitochondria in living cells

For Airy-SIM imaging: The COS-7 cells were labelled with PKmito RED (Nanjing Genvivo Biotech Co., Ltd. PKMR-1, CY-SC052)/PKmito DEEP RED (Cytoskeleton, CY-SC055) 15 minutes in DMEM. After labeling, wash the dye 2-3 times with new pre-warmed DMEM before imaging.

For LiveSR imaging: The U2OS cells were labeled with 250 nM MitoTracker Green FM (Invitrogen, M7514) 30 min before imaging.

For STED imaging: The COS-7 cells were labeled with IMMBright 660^58^ at 37°C for 10 min before imaging.

#### Labeling tubulin in living cells

For Airy-SIM imaging: We use SiR Tubulin Kit (Cytoskeleton, CY-SC002) to label tubulin in live COS-7 cells under the conconcentration of 1μM. Incubate the cells in the incubator with 5% CO^2^ at 37°C for 1 hour before imaging.

For LiveSR imaging: The tubulin-GFP plasmid was transfected into U2OS cells with Lipofectamine 3000 (Invitrogen, L3000) under the standard protocol.

#### Labeling lysosome in living cells

We use LysoView™ 488 (Biotium,70067) to stain the lysosome in COS-7 cells for 15-30 minutes without washing.

### Pearson correlation coefficient

Pearson correlation coefficient measures the similarity between two images:

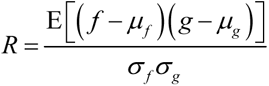

where *f* and *g* are two images to be compared, *μf* and *μg* are their average value, *σf,* and *σg* are their standard deviation value.

In this work, we use Pearson correlation to measure the correlation of the raw image with the noisy image and MRA deconvolved image in the simulation presented in Fig. 1 and Extended Data Fig. 2. Moreover, we also use the Pearson correlation to quantify the correlation between two organelles. To calculate the Pearson correlation between lysosome and tubulin displayed in Fig. 4c, we first segment the lysosome image, and calculate the Pearson correlation between the signal region in lysosome image with the microtubule image. Similarly, we subtract the mitochondrial distal end region and calculate its Pearson correlation with the ER image displayed in Fig. 5f.

### Mander’s overlap coefficient

Mander’s overlap coefficient (MOC) is another index that measures the correlation between two organelles. Compared with Pearson correlation, MOC has better interpretability and focuses on absolute colocalization. Here we also use MOC to cross-validate the colocalization of organelles. MOC is calculated as follows:

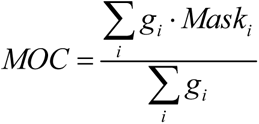

where *g* is the gray value of one organelle, and *Mask* is the binarized image of another organelle.

We calculate the MOC of lysosome and tubulin displayed in Fig. 4c by generating a *Mask* using a segmented lysosome image.

### Image decorrelation analysis

We use image decorrelation analysis^45^ to estimate the image resolution and evaluate image SNR from the view of frequency domain. After standard edge apodization that mitigates high-frequency artifacts, it calculates the cross-correlation of the image spectrum and its normalized spectrum. Then this process is repeated with the normalized spectrum additionally filtered by a binary mask. The decorrelation curve is expressed as follows:

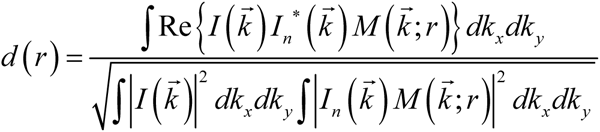

where 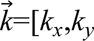 is the frequency-domain coordinate, *I* 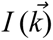 and *I_n_* 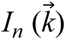 are the image Fourier spectrum and normalized spectrum, respectively, 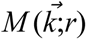 is the binary mask with a radius of *r*.

The normalized Fourier spectrum balances the contribution of signal and noise, which is crucial to differ them. As most information is recorded in the low-frequency region due to the band-limited nature of the optical system, the decorrelation curve would firstly increase with the increase of radius *r* until most signals are included, then it would decrease with increasing radius because noise contribution is larger. The maximum value of the decorrelation value *A*0 (0∼1) reflects the SNR metrics, which are used to evaluate the image SNR From the view of the Fourier domain.

### NanoJ-SQUIRREL resolution scaled error

The QUIRREL algorithm^46^ is employed to evaluate the resolution-scaled error of a resolution-enhanced image *f* based on low-resolution image *g* and resolution-scaling function (RSF). It starts by correcting the lateral mismatch of the two images, then find the optimal parameter to convolve the resolution-enhanced image *f* back to *g*:

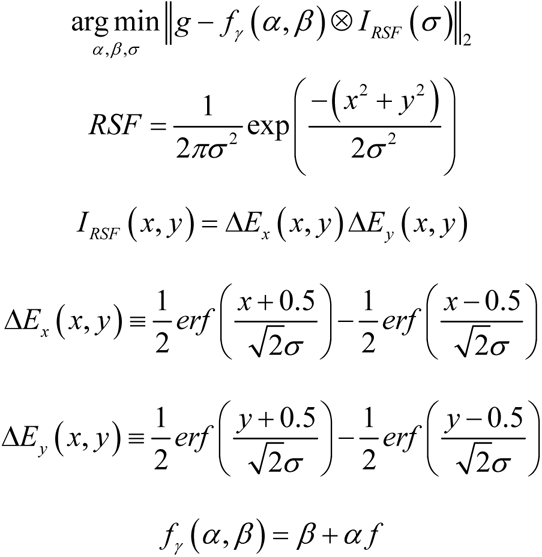

Then *f* is convolved back using the optimal parameter: *f*RS=*αf*+*β*, which is used to calculate the resolution-scaled error (RSE) and error map:

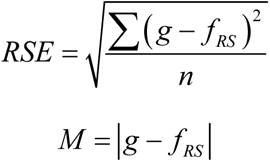

### SSIM

We use SSIM to measure the similarity between the convolved back image and the original image to evaluate the fidelity of the deconvolution algorithm as a supplement to the NanoJ-SQUIRREL analysis, which can consider the over-inferring artifacts. The SSIM between two images is calculated as follows:

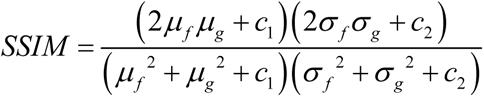

where *f* and *g* are two images to be compared, *μf* and *μg* are their average value, *σf,* and *σg* are their standard deviation value, *σf σg* is the covariance of *f* and *g*, *c*1 and *c*2 are two constants to stabilize the result (*c*1=(*k*1*L*)^2^, c2=(*k*2*L*)^2^, *L* is the dynamic range of pixel value, *k*1=0.01, *k*2=0.03).

### FWHM analysis

We employ FWHM analysis of the tubulin filament to quantitively assess the fidelity of the statistical and physical deconvolution method that displayed in Extended Data Fig. 3. The gray value across the microtubule filament is measured and interpolated to calculate the FWHM value on a finer sampling. We also determine the theoretical unblurred FWHM by generating Gaussian functions that have the same FWHM with the original image and SIM PSF, which is symbolled by *G_original_* and *G_PSF_*. We then calculate the Gaussian function of theoretically unblurred image *G_unblurred_* by finding the optimal *σ* value that satisfies *G_unblurred_*\**G_PSF_*= *G_original_*, which determines the FWHM in the theoretically unblurred image.

### Spectrum boundary-based resolution estimation

Besides decorrelation analysis resolution analysis, we also incorporate spectrum boundary determination to evaluate the image resolution in Fig. 1d and f, which is a more straightforward way to estimate the image resolution under severe noise. We first do Fourier transform of the original image, then accumulate the spectrum line by line to form a one-dimensional intensity curve. We then start from the maximum point and search for the point when the intensity is 1% of the maximum value, which is rendered as the boundary of the spectrum. The corresponding spatial resolution is calculated as:

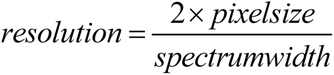

### SNR, PSNR, and SBR estimation

Considering that there is no GT image in practical imaging, we calculate the image SNR (dB) using the following commonly used formula:

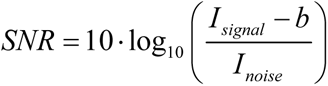

where *I_signal_* denotes the intensity of the signal, *I_noise_* denotes the intensity of noise, and *b* denotes the background intensity.

We estimate *I_signal_* by calculating the average intensity in a selected region which is taken as the signal, and estimate *I_noise_* and *b* by calculating the intensity standard deviation and average value in a selected non-signal region.

The PSNR is calculated as follow:

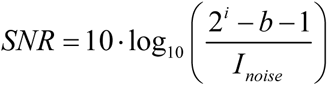

where *i* is the bit of the image.

A similar approach is used to calculate SBR:

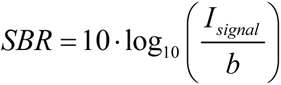

## Author Contribution

Y. Hou and P. Xi conceived the idea of multi-resolution analysis-based fluorescence deconvolution algorithm. P. Xi supervised the research. Y. Hou fabricated the mathematical framework, wrote the code and composed all the figures and videos. M. Li and Y. Hou devised the biological experiments. W. Wang, Y. Fu and X. Ge conducted the biological experiments. Y. Hou, P. Xi and M. Li wrote the manuscript with inputs from all the authors. All authors discussed the results presented in the manuscript.

## Acknowledgements

This work was supported by the National Key R&D Program of China (2022YFC3401100), and the National Natural Science Foundation of China (62025501, 31971376, 92150301). We thank National Center for Protein Sciences at Peking University in Beijing, China, for assistance with LiveSR, SoRa, GE DeltaVision OMX SIM, and Abberior STED-Facility super-resolution imaging. We thank Prof. Jingyan Fu at China Agricultural University for providing the centrosome sample.

---

**Extended Data Fig. 1.**
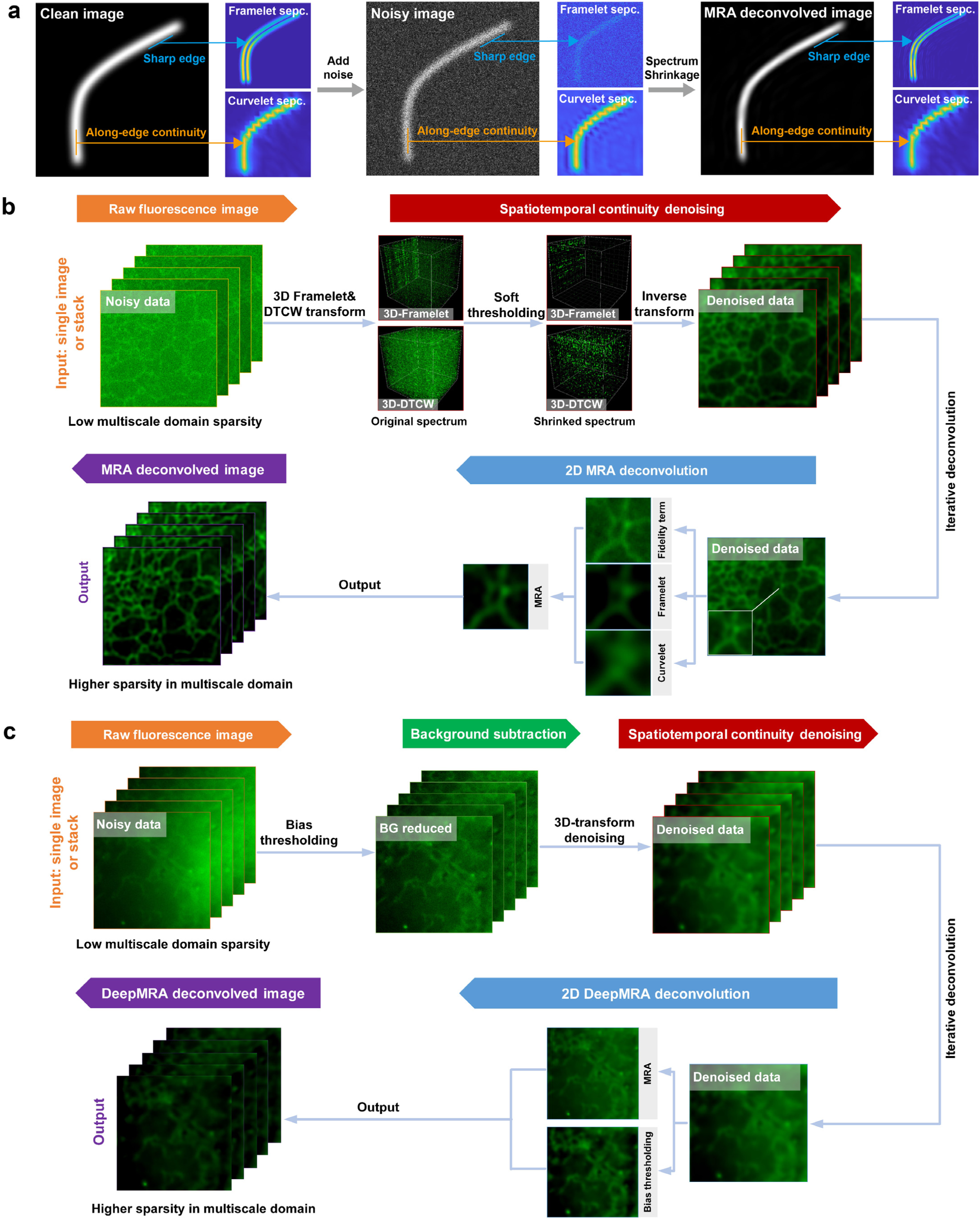
Schematic diagram of the notion and procedure for MRA and DeepMRA deconvolution. **a.** Principle of framelet and curvelet denoising. **b.** Flow chart of MRA deconvolution algorithm, including two major processing steps: spatiotemporal continuity denoising (optional), and 2D MRA deconvolution. **c.** Flow chart of DeepMRA deconvolution algorithm, including three major processing steps: background subtraction (optional), spatiotemporal continuity denoising (optional), and 2D DeepMRA deconvolution.

**Extended Data Fig. 2.**
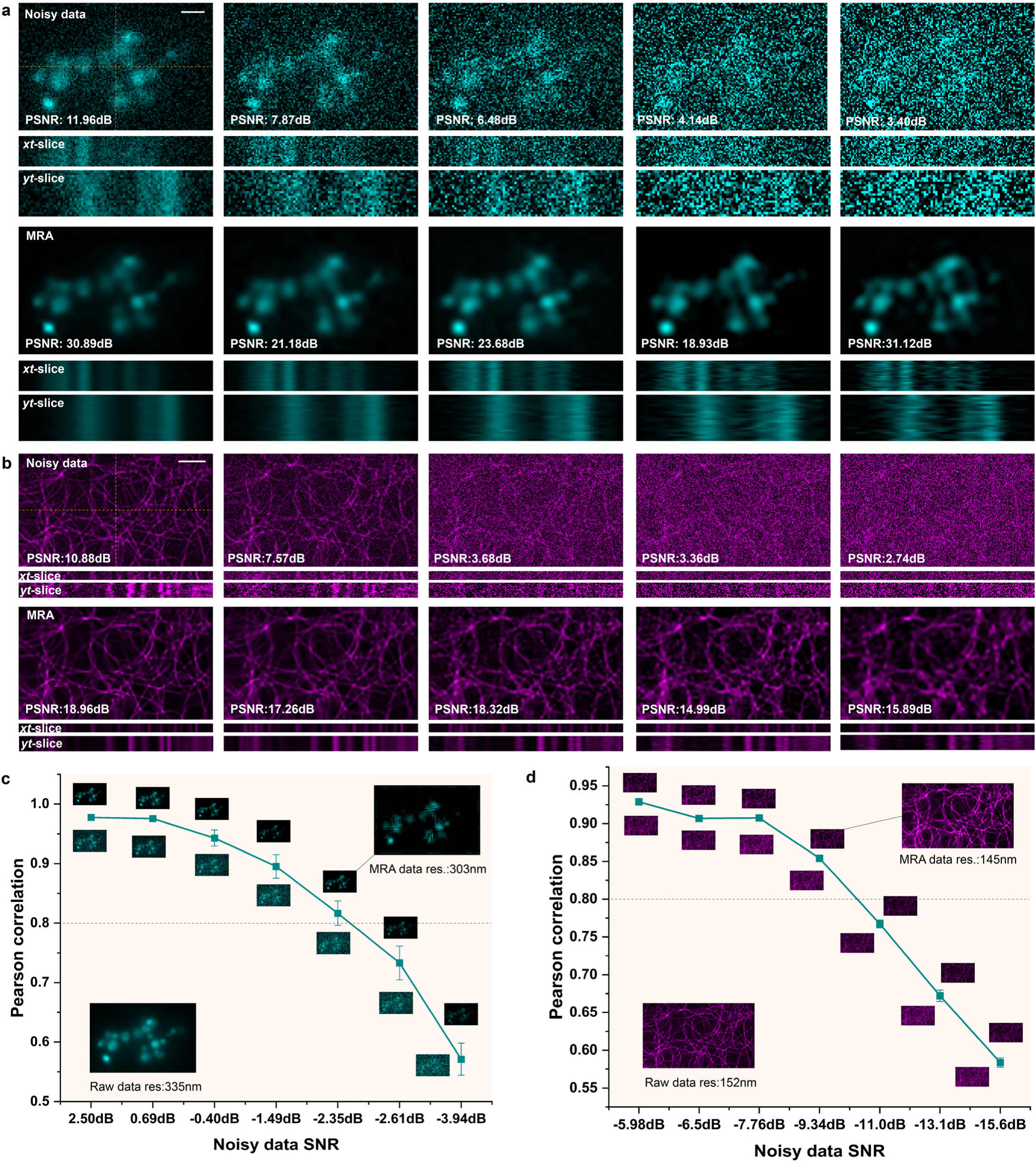
MRA deconvolution resolves time-lapse images from severe noise contamination. Lysosome **a** and microtubule **b** time-lapse images with varying Gaussian noise levels, and the corresponding MRA deconvolution results. Pearson correlation of resolution-maintaining MRA deconvolution of lysosome **c** and tubulin **d** images with different Gaussian noise levels. Scale bars: 1 *μ*m (a), 2 *μ*m (b).

**Extended Data Fig. 3.**
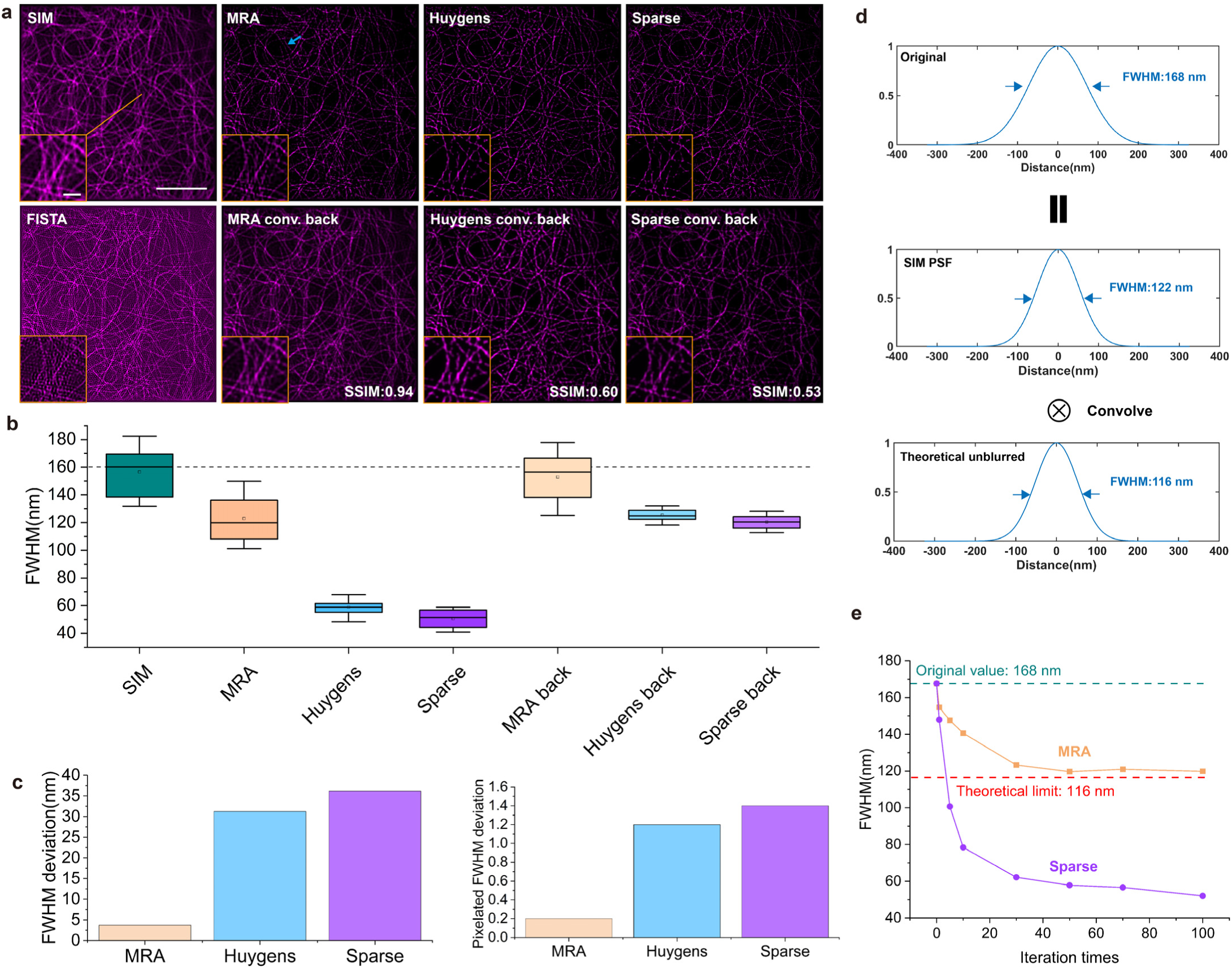
Quantitative assessment of the fidelity of physical and statistical deconvolution. **a.** SIM microtubule image, and the FISTA, MRA, Huygens, and Sparse deconvolution results, and the corresponding convolved back results. The SSIM value is calculated using the raw SIM image as a reference. **b.** Box chart of the FWHM of tubulin filament in each graph (*n*=10). **c.** The average value of the FWHM deviation of the MRA, Huygens, and Sparse convolved back results. The pixelated FWHM deviation is calculated by discretizing the FWHM value first. **d.** Determination of theoretically unblurred FWHM of the blue arrow denoted filament in a. **e.** Variation of the FWHM as the change of iteration times in MRA and the RL iteration times in Sparse. Scale bars: 5 *μ*m (a), 1 *μ*m (a subgraph).

**Extended Data Fig. 4.**
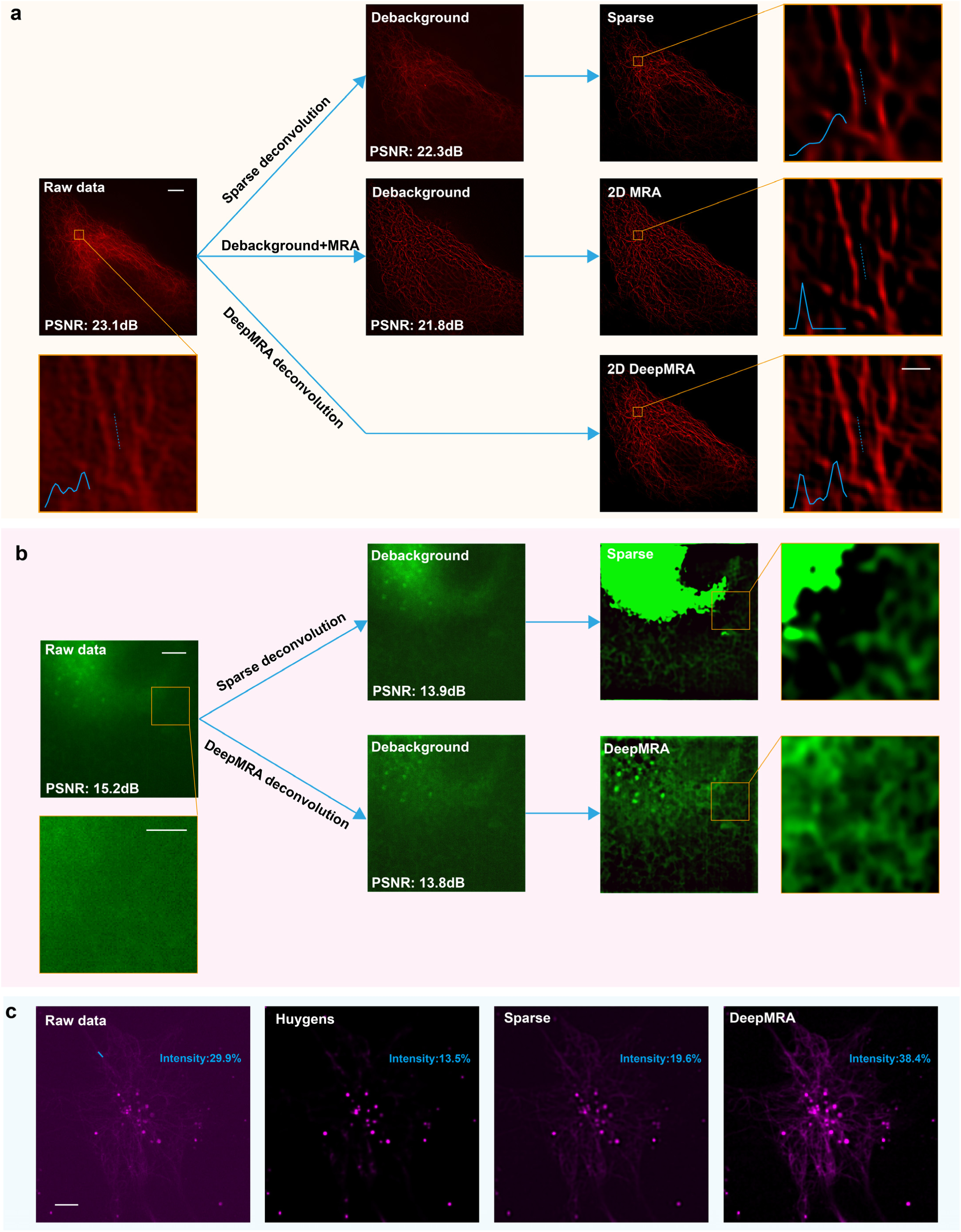
Comparison of different schemes of background-mitigated deconvolution in fluorescence images. **a.** Deconvolution of microtubule image using Sparse, MRA after debackground, and Deep MRA. **b.** Deconvolution of ER image using Sparse, MRA after debackground, and DeepMRA. **c.** Deconvolution of microtubule image with ultra-bright point using Huygens, Sparse, and DeppMRA. Raito denotes the intensity ratio of the red line average to the maximum. Scale bars: 5 *μ*m (a Left, b Left, c), 2 *μ*m (b Right), 0.5 *μ*m (a Right).

**Extended Data Fig. 5.**
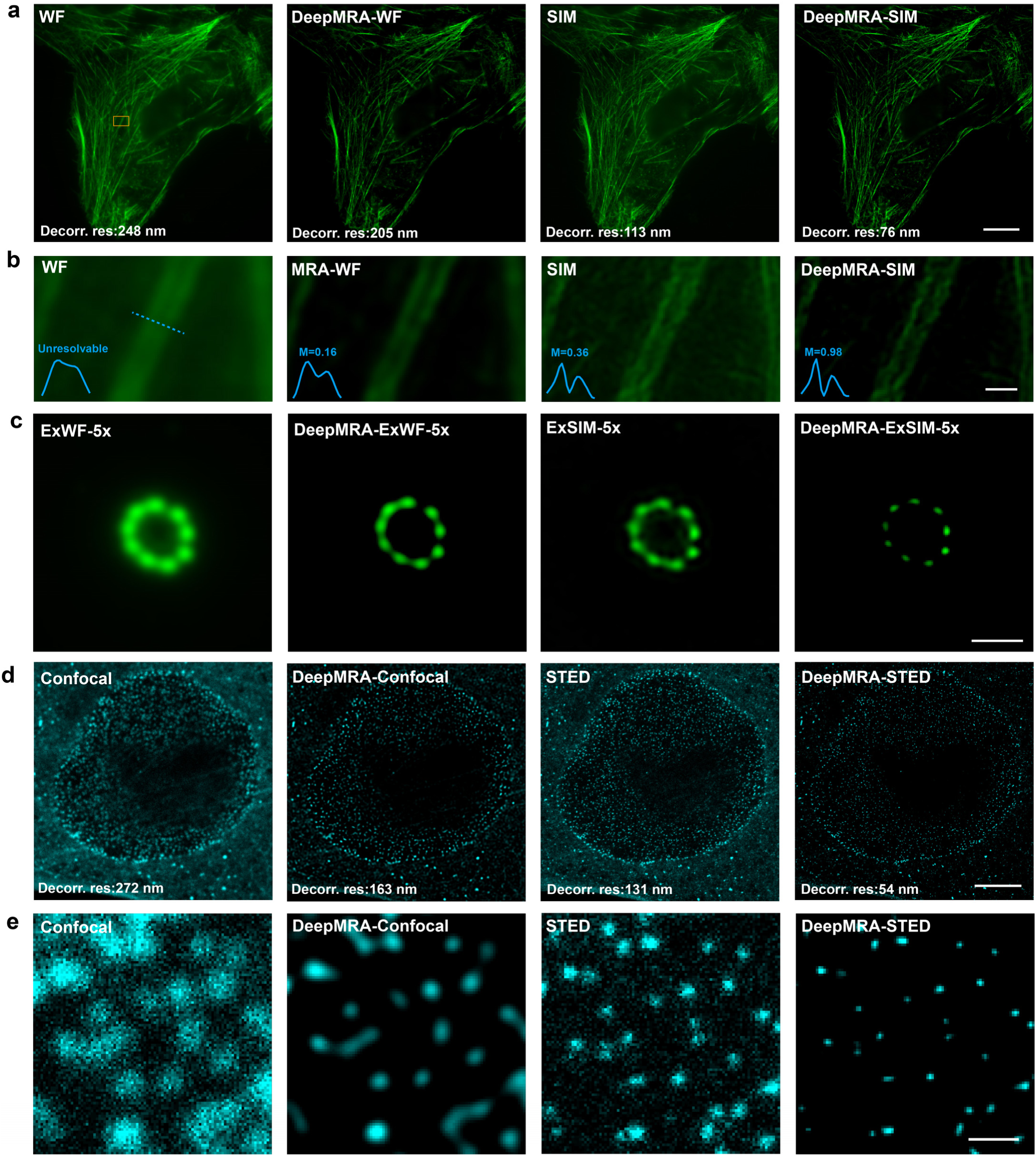
DeepMRA deconvolution improves fluorescence image resolution. **a.** U2OS cell actin image captured by wide-field microscopy and SIM, and the corresponding DeepMRA deconvolution results. **b.** Magnified yellow boxed region in a. The intensity profile of the dotted line and its modulation factor is shown at the left bottom of each graph. **c.** Expansion microscopy image of the centriole, and the corresponding DeepMRA deconvolution results. **d.** Mouse embryonic fibroblasts nuclear pore image captured by confocal and STED microscopy^11^, and the corresponding DeepMRA deconvolution results. **e.** Magnified orange boxed region in d. Scale bars: 10 *μ*m (a), 5 *μ*m (d), 0.5 *μ*m (b, e), 0.2 *μ*m (c).

**Extended Data Fig. 6.**
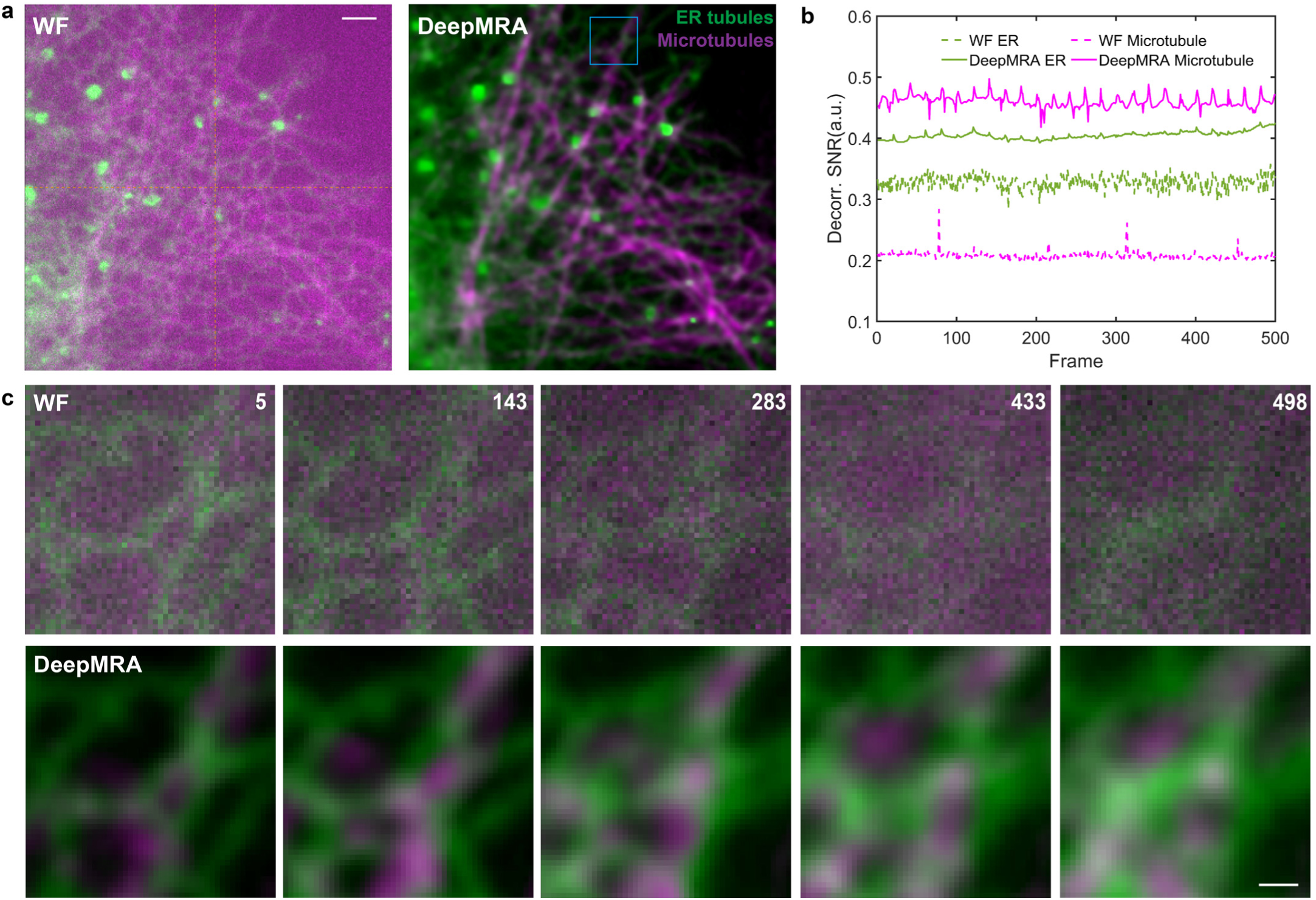
DeepMRA deconvolution assists imaging of ER tubules (green) and microtubules (magenta). **a.** The wide-field time-lapse images of co-labeled ER tubules and microtubules, and the DeepMRA deconvolution results. The imaging lasts 5 minutes and 33 seconds with 500 frames. **b.** SNR was obtained by decorrelation analysis at different frames. **c.** Magnified blue boxed region in a. Scale bars: 2 *μ*m (a), 0.5 *μ*m (c).

**Extended Data Fig. 7.**
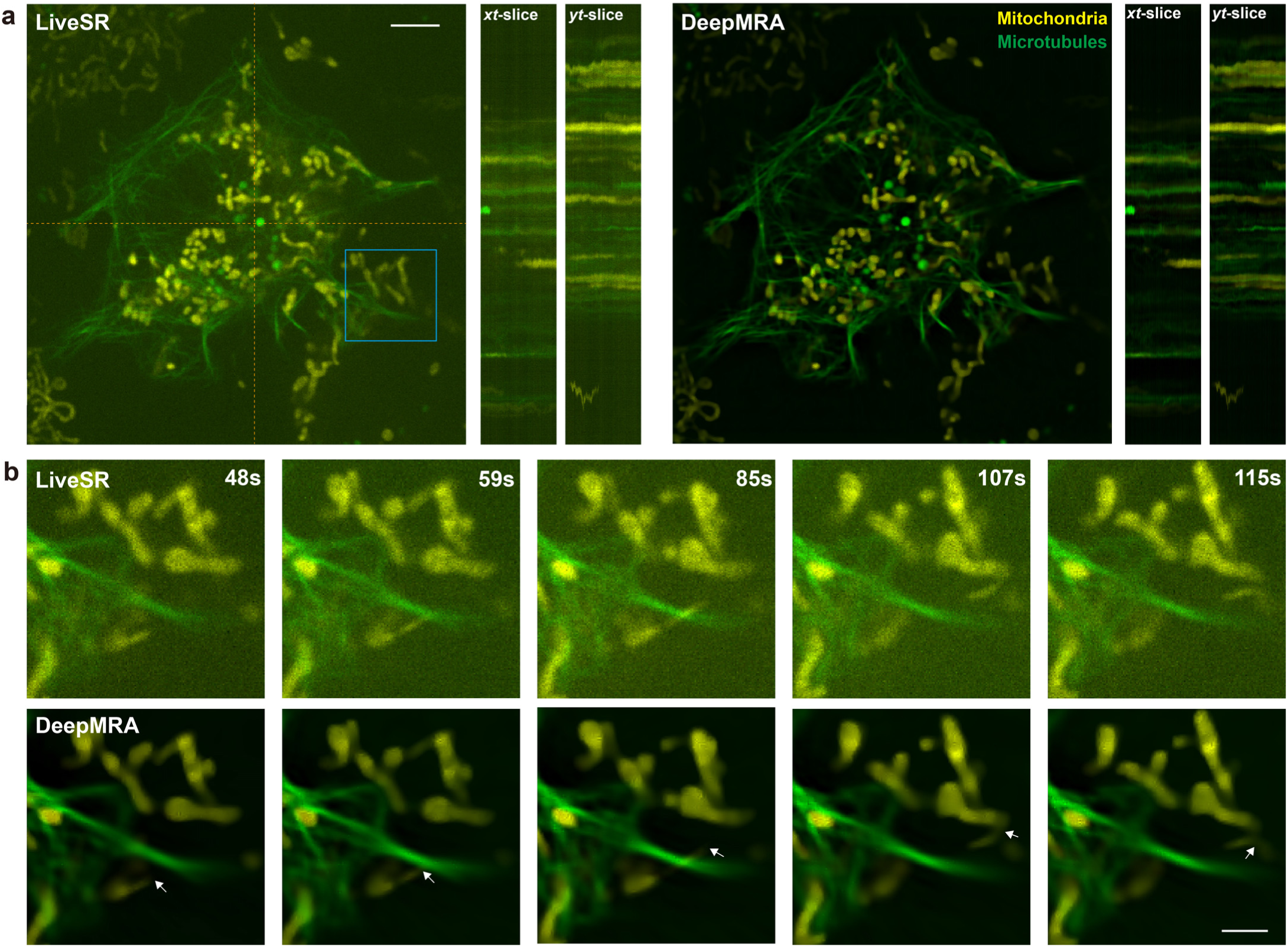
DeepMRA deconvolution assists imaging of mitochondria and microtubules interaction. **a.** The LiveSR time-lapse images of co-labeled mitochondria and microtubules in U2OS cell, and the DeepMRA deconvolution result. It consists of 120 frames with a duration of 120 s. **b.** Magnified blue boxed region in a. Scale bars: 5 *μ*m (a), 2 *μ*m (b).

**Extended Data Fig. 8.**
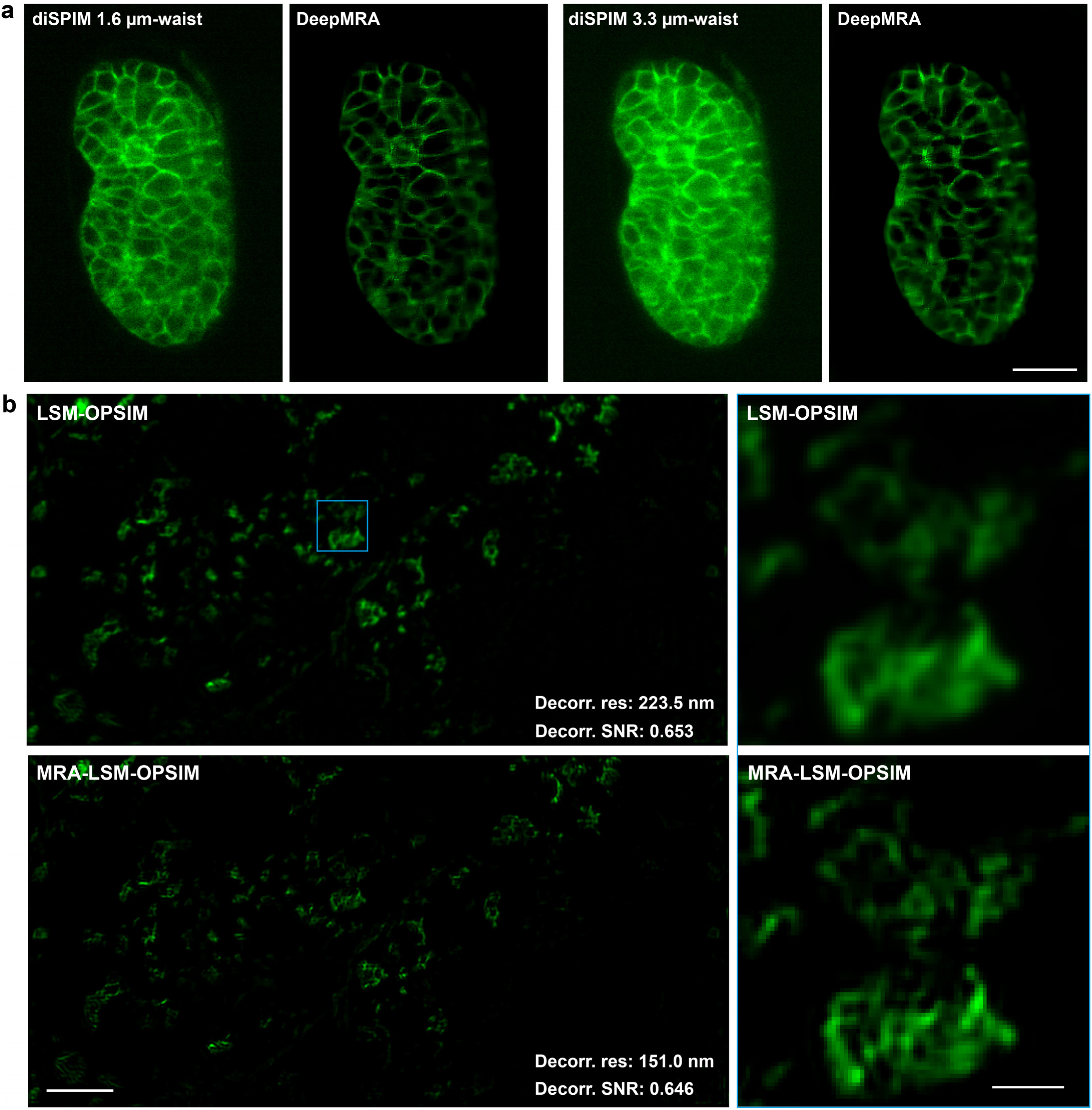
DeepMRA assists light-sheet microscopy (LSM) imaging. **a.** diSPIM image of C. elegans embryos with 1.6 and 3.3 *μ*m Gaussian beam wrist^59^, and the corresponding DeepMRA deconvolution result. **b.** OPSIM image of spinal cord slice^60^ and the MRA deconvolution result. Scale bars: 10 *μ*m (a), 5 *μ*m (b Left), 1 *μ*m (b Right).

